# Using regions of homozygosity to evaluate the use of dogs as preclinical models in human drug development

**DOI:** 10.1101/2020.01.08.898916

**Authors:** SP. Smieszek, MP. Polymeropoulos

## Abstract

Animals are used as preclinical models for human diseases in drug development. Dogs, especially, are used in preclinical research to support the clinical safety evaluations during drug development. Comparisons of patterns of regions of homozygosity (ROH) and phenotypes between dog and human are not well known. We conducted a genome-wide homozygosity analysis (GWHA) in the human and the dog genomes.

We calculated ROH patterns across distinct human cohorts including the Amish, the 1000 genomes, Wellderly, Vanda 1k genomes, and Alzheimer’s cohort. The Amish provided a large cohort of extended kinships allowing for in depth family oriented analyses. The remaining human cohorts served as statistical references. We then calculated ROH across different dog breeds with emphasis on the beagle - the preferred breed used in drug development.

Out of five studied human cohorts we reported the highest mean ROH in the Amish population. We calculated the extent of the genome covered by ROH (F_ROH_) (human 3.2Gb, dog 2.5Gb). Overall F_ROH_ differed significantly between the Amish and the 1000 genomes, and between the human and the beagle genomes. The mean F_ROH_ per 1Mb was ∼16kb for Amish, ∼0.6kb for Vanda 1k, and ∼128kb for beagles. This result demonstrated the highest degree of inbreeding in beagles, far above that of the Amish, one of the most inbred human populations.

ROH can contribute to inbreeding depression if they contain deleterious variants that are fully or partially recessive. The differences in ROH characteristics between human and dog genomes question the applicability of dog models in preclinical research, especially when the goal is to gauge the subtle effects on the organism’s physiology produced by candidate therapeutic agents. Importantly, there are huge differences in a subset of ADME genes, specifically cytochrome P450 family (CYPs), constituting major enzymes involved in drug metabolism. We should hesitate to generalize from dog to human, even if human and beagle are relatively close species phylogenetically

## INTRODUCTION

Animals are used as preclinical model for human diseases in drug development. While central to drug development, there is lack of validity for many of such tests. Evidential weight gain accrued by the use of animal data was assessed in a recent study, evaluating the probability that a certain new tested drug may prove toxic to humans^1^. The authors calculated likelihood ratios (LR) for a large set of drugs. They have shown that lack of toxicity in fact provides no weight gain for lack of adverse drug reactions, as well as large inconsistencies in LR when predicting from animals to human^1^.

Dogs, especially, are used in preclinical research to support the clinical safety evaluations during drug development, however, dogs are a particularly inbred species. Recently, EMBARK project conducted large scale characterization for the dog ROH density maps for 2500 dogs comparing a range of breeds^2^. The presented results recorded ROH down to 500 kilobases. Authors report more than ‘678 homozygous deleterious recessive genotypes in the panel across 29 loci, 90% of which overlapped with ROH’^2^. The study constitutes one of the most comprehensive evaluations of dog genome across several breeds of dogs, showing great degree of variation among breeds. Patterns of ROH in dogs have seem to imply a high degree on inbreeding across multiple breeds of dogs, regions furthermore associating with deleterious variants^2^. Long ROH are enriched in deleterious variation^3^. It has been further shown that inbreeding as observed via genomic measures reduces fecundity in Golden Retrievers^4^. It is well established that high degree of inbreeding results in decreased reproductive fitness as discussed initially by Charles Darwin in his initial observations in plants followed by in humans^5^,^6^. It is hence well established that inbreeding increases the incidence of recessive disease as discussed already in 1902 in the case of alkaptonuria^7^ and in cases of Mendelian disorders such as Tay-Sachs^8^. Moreover, the implications of extensive inbreeding were recently demonstrated and quantified in a sequencing study of grey wolves of Isle Royale, where inbreeding depression which has brought wolves to the brink of extinction likely an effect accounted for increased homozygosity of the deleterious variants^9^. The comparison of regions of homozygosity (ROH) patterns between dog and human has not been characterized. In the present study we focus on exactly that comparison. A ROH is defined as a continuous stretch of DNA sequence without heterozygosity in the diploid state (min ROH >= 1.5 Mb). We calculated ROH patterns across distinct human cohorts: the Amish, IGSR 1000 Genomes^10^, Wellderly^11^, Vanda 1K genomes using both microarray and sequencing

We have incorporated several datasets in the analysis to have a broader view upon the potential differences in and between the two species as across breeds, across human subpopulations. The Amish with large families, well documented family history accessible via Anabaptist Genealogy Database constitute a valuable resource for genetic studies^12^. Amish constitute an example of a highly inbred population, in fact ROH were first described in the Amish. The 1000 genomes serve as mixed control representation of individuals. Vanda 1000 genomes is a control set of whole genome sequencing samples with extensive phenotyping. The Wellderly present a well-being well aged cohort of mixed ethnicity. The Alzheimer’s cohort is a mixed ethnicity cohort offering yet another point of comparison. Using the same settings across datasets we calculated ROH across different dog breeds (EMBARK project) with emphasis on the beagle, as it is the preferred breed in drug development. We evaluate the degree of inbreeding in dogs compared to human populations. We furthermore focus on the CYP family, as they are the major class of enzymes involved in drug metabolism, accounting for 75% of the total metabolism^13^. The results should allow one to investigate the genetic validity of dogs as preclinical models for human drug development.

## RESULTS

A ROH is defined as a continuous stretch of DNA sequence without heterozygosity in the diploid state (min ROH = established usually between 500kb and 1.5 Mb).We estimate ROH using PLINK^14^. The hypothesis is that that deleterious genotypes will be enriched in the ROH regions as compared by background. We characterize the distribution of ROH across dog breeds, across human population cohorts as well as between human and the dog evaluated on the same parameters. **Figure 1** displays the length and the number of ROHs identified across numerous dog breeds.

**Figure 1.**
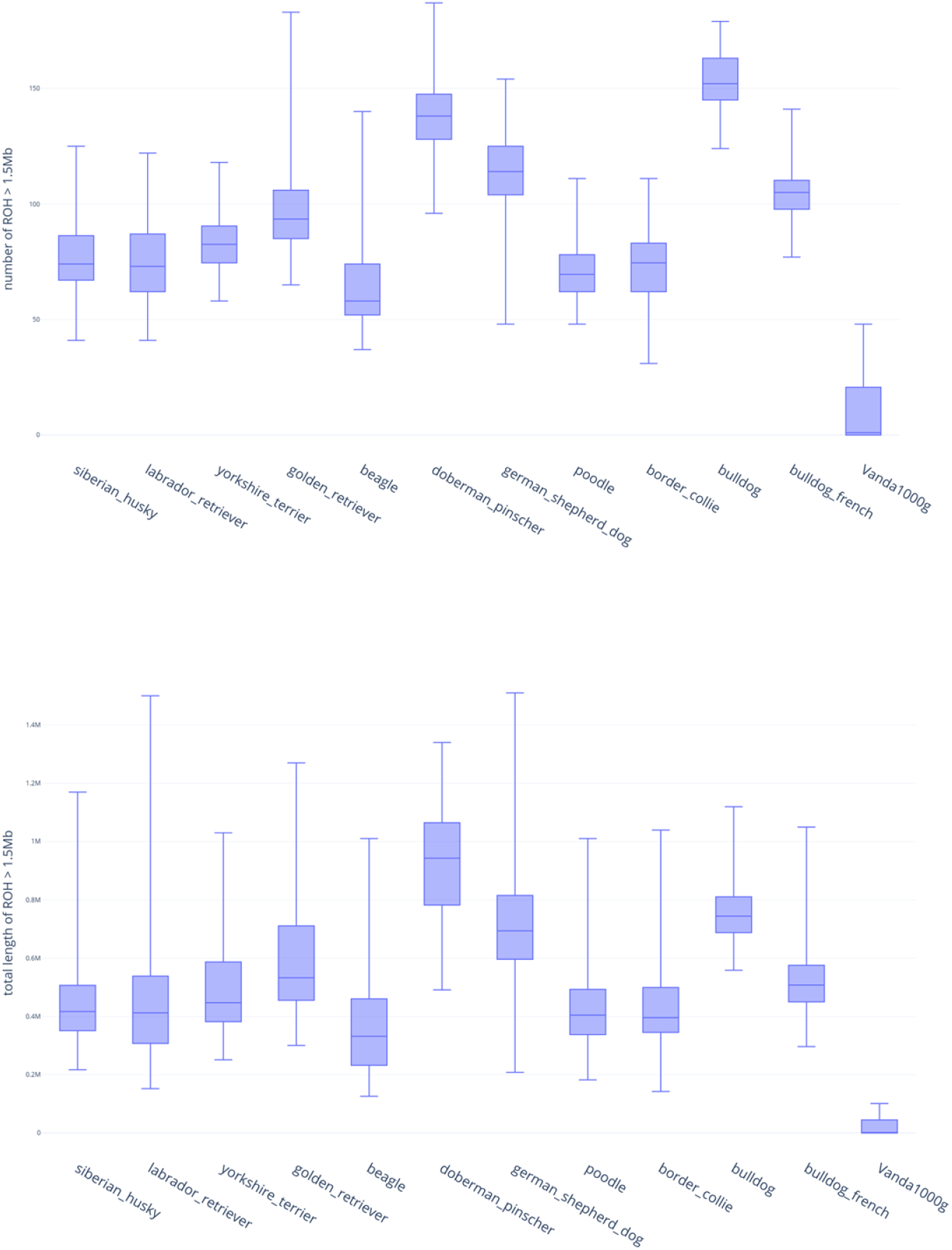
The total number and length of ROH identified across dog breeds as compared to mixed ethnicity human cohort: Vanda 1000 genomes.

Vanda 1k and the Wellderly ROH analysis was conducted on whole genome sequencing data. When we considered both the location and the allelic form of the ROHs, we were able to separate the populations by PCA, demonstrating that ROHs contain information on the demographic history and structure of a population. We calculated the extent of the genome covered by ROH (F_ROH_) (human 3.2Gb, dog 2.5Gb) as displayed in **Table 1**.

**Table 1.**
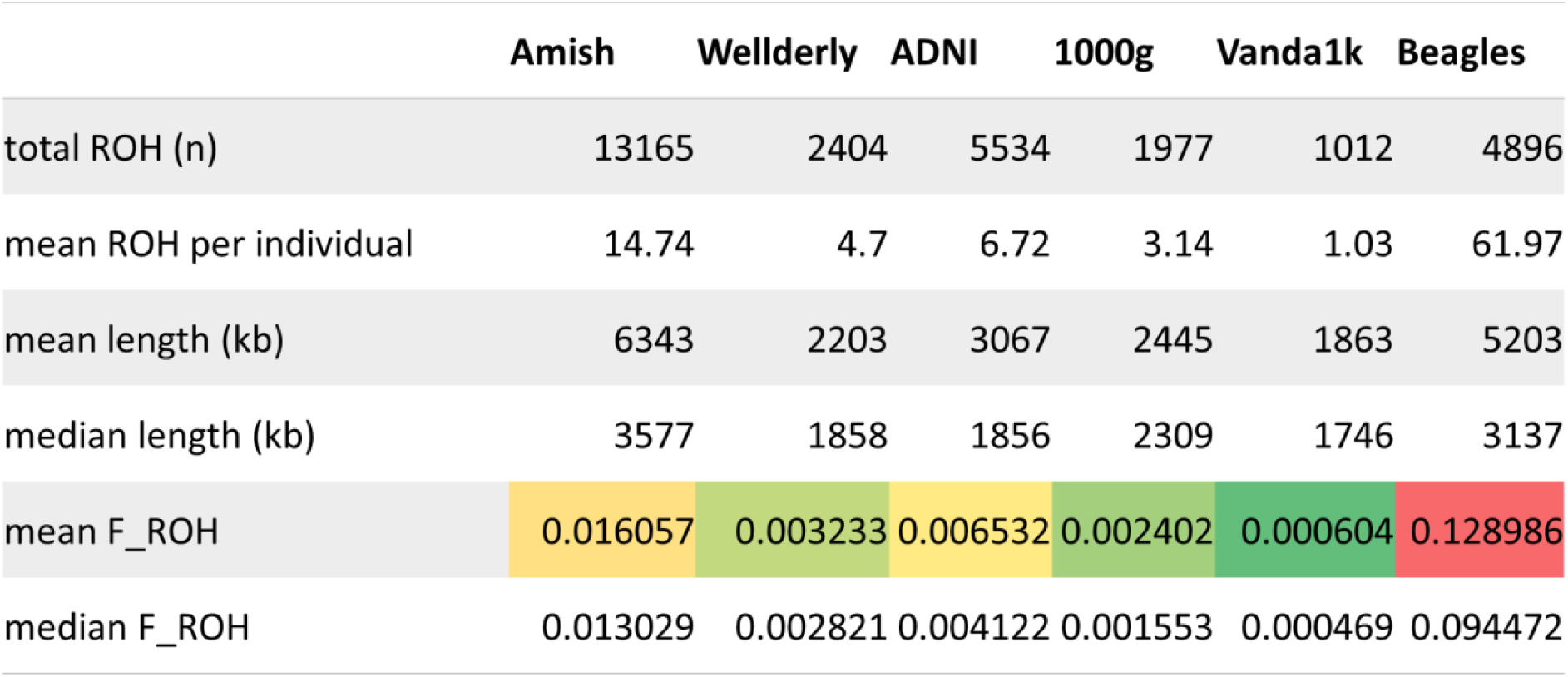
Displays the total, mean ROH and (F_ROH_) across human cohorts compared to dogs.

In the Amish population, one of the most inbred human populations, the F_ROH_ was 12x smaller in comparison to the beagle population. F_ROH_ differed significantly between the Amish and the 1000 genomes, and between the human and the beagle genomes. The mean F_ROH_ per 1Mb was ∼16kb for Amish, ∼0.6kb for Vanda 1k, and ∼128kb for beagles. This result demonstrated the highest degree of inbreeding in beagles, far above that of the Amish, one of the most inbred human populations also displayed on **Figure 2**. Figure 2 exemplifies the significantly higher observed ROH as compared to several human populations including the Amish. **Figure 3** further compares the human populations and the specific regions of high ROH across the human genome.

**Figure 2.**
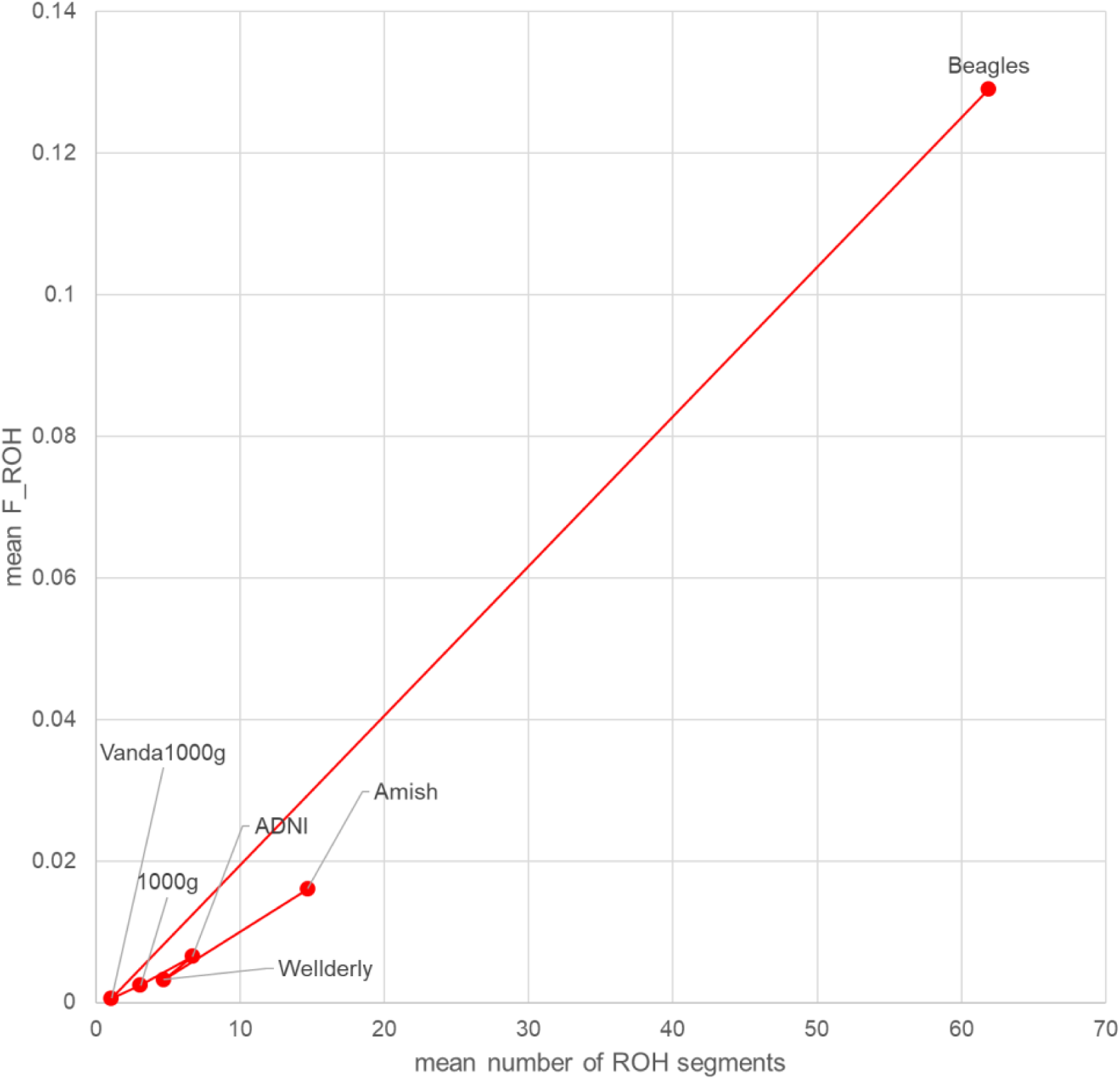
Mean F_ROH on the y axis and mean number of ROH segements across several human and dog cohorts.

**Figure 3.**
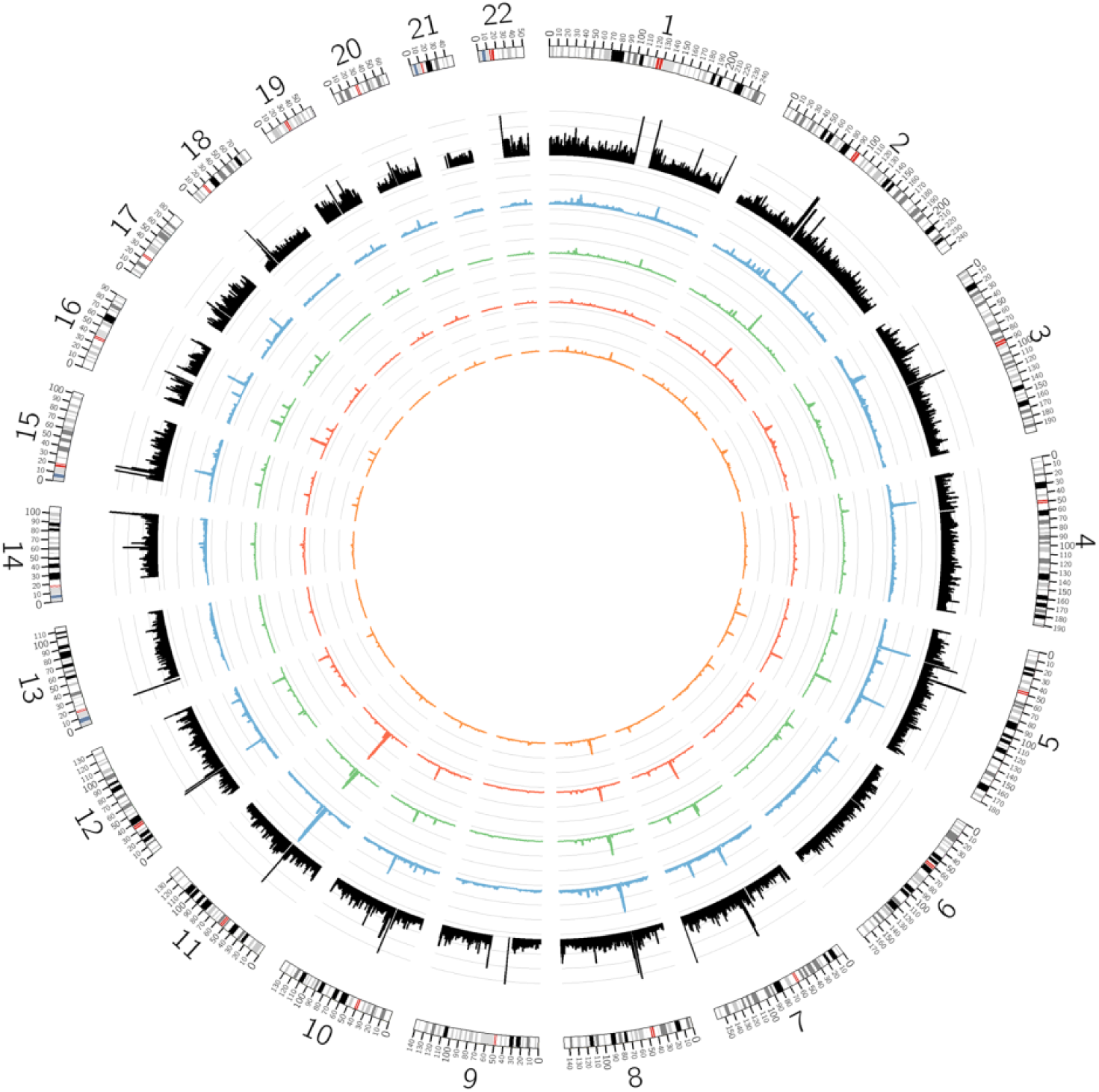
ROH patterns across human cohorts: Human genome arranged circularly end-to-end: from inside to outside, the rings display ROH across different human cohorts starting from the innermost: 1000g, ADNI, Wellderly and Amish (blue). The outermost ring represents conservation score averaged per region of the human genome.

Currently, we classify 298 genes that encode Phase I and II drug metabolizing enzymes, transporters, and modifiers as ADME genes, with CYPs constituting a major family. To further evaluate the consequences we focused on CYP family including most common CYPs involved in metabolism of human drugs including CYP2D6, CYP2B’s, CYP2C’s, CYP3A’s. We mapped orthologues between dogs and human using BIOMART^15^ (orthologues, from human to dogs). We report that beagles alone, F_ROH, would be 25% so even higher than our average in beagles, 12.9%. For humans that estimate would be 1/1000 on the same parameter. The results are presented in **Figure 4**.

**Figure 4.**
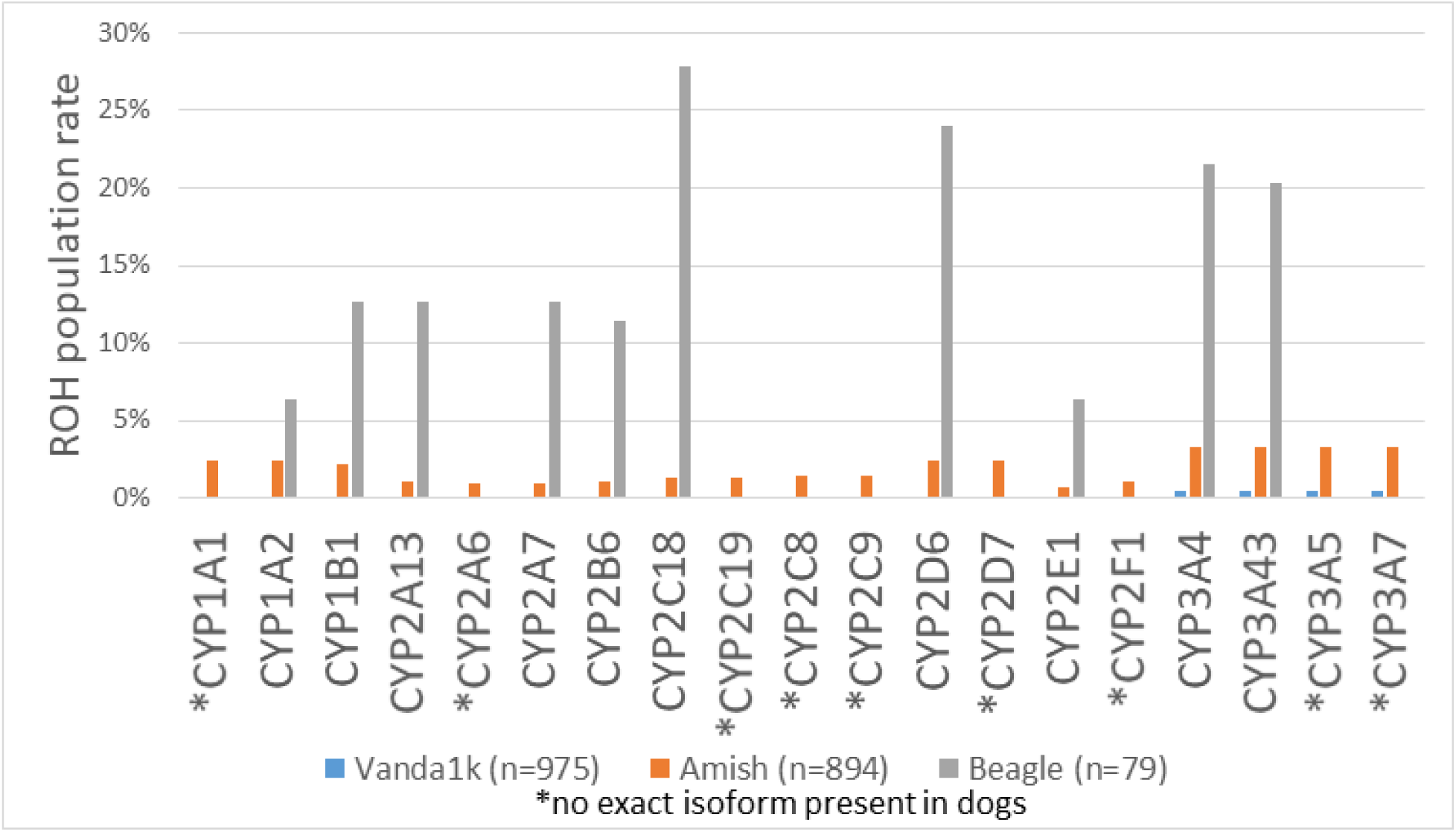
ROH patterns in CYPs compared between humans and dogs. Especially high signal is seen for CYP2C18, CYP2D6, CYP3A4, CYP3A43 amongst others.

## DISCUSSION

The study found a high degree of homozygosity across the genomes of beagle dogs, an indication of high degree of inbreeding. High degree of inbreeding creates an evolutionary bottleneck and exposing homozygous deleterious variants in the genome of the species. Furthermore due to heterozygote advantage as outlined by overdominance hypothesis, the long term loss of diversity furthermore reduced heterozygote advantage. For example, the risk for autosomal recessive disease is proportional to the degree of parental relationship in in consanguineous families’^16^. In humans total length of ROH per individual show considerable variation across individuals and populations with higher values occurring more often in populations with known high frequencies of consanguineous unions. Even across the genome the distribution is far from uniform^17^. Across the genome the variation is correlated with recombination rate, as well as with signals of recent positive selection. Importantly, long ROH are more frequent in genomic regions harboring genes associated with autosomal-dominant diseases^18^. Interestingly the top-ranked ROH hotspot in the human genome as assessed by the authors is located on chromosome 2p in a region with low recombination rates and containing CYP26B1^18^.

We compared the degree of genetic inbreeding in beagle dogs using the measure of runs of homozygosity (ROH). This therefore focus on contiguous regions of the genome that are inherited as identical from both parents. The study showed that unrelated humans show these regions of homozygosity in about 1/1000 of their genomes while beagle dogs do so in 12.8 percent of their genome. In other words beagle dogs are on average 100 times more inbred than humans. In the same study researchers examined the degree of inbreeding in the genetic isolate of the Amish in Pennsylvania which showed that they were about 12 times more inbred than the general population. It is well accepted that drug efficacy and safety studies in genetic isolates such as the Amish cannot be generalized for the population at large.

The ADME genes constitutes a highly polymorphic set of genes involved in Absorption, Distribution, Metabolism, and Excretion of drugs. This was exemplified in a recent effort to characterize the distribution of 95 polymorphisms in 31 core ADME genes in 20 populations worldwide^19^. The present effort to evaluate CYPs not within but in the context of between species as displayed on Figure 4, comes without a surprise. CYP2D6, another gene with allelic variants and encoding enzymes with variable degrees of activity, is associated with the development of hepatotoxicity after use of certain pharmaceutical agents. The CYP2D6 gene is polymorphic with over 150 allelic variants as defined by (https://www.pharmvar.org/gene/CYP2D6)^20^. The high levels of variation in the frequencies of alleles of polymorphic pharmaco-genes between diverse ethnic groups may be responsible for severe adverse reactions to as well as altered potentially efficacy of a wide variety of drugs^21^.

Beagle dogs are routinely used for studying the safety of new pharmaceuticals. This practice has been in place for a hundred years and followed by pharmaceutical developers and regulators. The present study is significant in that it casts a big question over the predictive validity of these commonly conducted dog studies. The very high degree of inbreeding seen would suggest that beagle dogs would be a poor model to predict toxicity even for other dog breeds and of course an inappropriate model to do so for humans. Reliance upon the beagle dog to predict human drug safety is not warranted especially if metabolism of a drug is known to be driven by CYP family in example. The results of this study should lead to the abandonment of the routine use of dogs in human drug toxicology studies and urge researchers and regulators to instead adopt appropriate and relevant scientific approaches in order to ensure human drug safety.

## CONLCUSION

The fraction of the genome covered by ROH (F_ROH_) was significantly different between human populations (Amish and 1000 genomes) and between humans and beagles. In the Amish population, one of the most inbred human populations, the F_ROH_ was 12 times smaller in comparison to the beagle population. ROH patterns in beagles increase their susceptibility to inbreeding depression and may introduce deleterious or recessive traits into their genome.Due to the high degree of inbreeding observed in beagles, preclinical research should use caution when generalizing from dog to human (especially when the enzymes metabolizing the drug are known), despite the physiological similarities between the species.

## METHODS

### Datasets

### ROH detection

#### Genomic data analysis ROH

Calculation of ROH in human cohorts including the Amish, IGSR 1000 genomes, Wellderly, Vanda 1k genomes. We detected ROH using PLINK. ROH scores reflect the probability of a stretch of SNPs being homozygous due to LOH and they are determined via the homozygosity frequency for each SNP in the genome. We detected ROH using PLINK on the EMBARK dog data. We next calculated the extent of the genome covered by ROH (F_ROH_). ROH were defined as runs of at least 50 consecutive homozygous SNPs spanning at least 1500 kb, with less than a 1000 kb gap between adjacent ROH and a density of SNP coverage within the ROH of no more than 50 kb/SNP, with one heterozygote and 5 no calls allowed per window. Detection of ROH, depletion with confidence scores, LD in ROH, region analysis and comparison across species

### DNA quantification

Incoming nucleic acid samples are quantified using fluorescent-based assays (PicoGreen) to accurately determine whether sufficient material is available for library preparation and sequencing.

### DNA integrity

DNA sample size distributions are profiled by a Fragment Analyzer (Advanced Analytics) or BioAnalyzer (Agilent Technologies), to assess sample quality and integrity.

### Genotyping

At the NYGC, we run the HumanCoreExome 24v1.3 array for all human DNA samples that we sequence.

### WGS library preparation and sequencing, Truseq PCR-free (450bp)

Whole genome sequencing (WGS) libraries were prepared using the Truseq DNA PCR-free Library Preparation Kit

#### WGS Germline analysis part I

Whole Genome data were processed on NYGC automated pipeline. Paired-end 150 bp reads were aligned to the GRCh37 human reference (BWA-MEM v0.7.8) and processed with GATK best-practices workflow (GATK v3.4.0).

The mean coverage is 35.8, it reflects the samples average. The coverage for the FLG region tracks with the coverage for the whole genome. The coverage information per FLG region per sample is provided in the supplementary file.

All high quality variants obtained from GATK were annotated for functional effects (intronic, intergenic, splicing, nonsynonymous, stopgain and frameshifts) based on RefSeq transcripts using Annovar (http://www.openbioinformatics.org/annovar/)^22^. Additionally, annovar was used to match general population frequencies from public databses (Exac, GnomAD, ESP6500, 1000g) and was used to prioritize rare, loss-of-function variants.

https://www.ncbi.nlm.nih.gov/pubmed/30452466

https://www.nature.com/articles/s41598-019-43610-y

https://www.frontiersin.org/articles/10.3389/fgene.2019.00007/full

https://www.fasebj.org/doi/abs/10.1096/fasebj.2019.33.1_supplement.814.10

https://www.ncbi.nlm.nih.gov/pmc/articles/PMC4013674/pdf/ijms-15-06990.pdf

https://www.ncbi.nlm.nih.gov/pmc/articles/PMC4548351/pdf/JCTH-3-099.pdf

https://journals.plos.org/ploscompbiol/article?id=10.1371/journal.pcbi.1005280

